# Reconfiguration of default mode metabolic demands and network organization during acute stress

**DOI:** 10.64898/2026.08.14.744771

**Authors:** Gabriel Schlosser, Christian Milz, Pia Falb, Samantha Graf, Sarah M Tüchler, Matej Murgaš, Alexandra Mayerweg, Clemens Schmidt, Ivan Pörnbacher, Lukas Artmeier, Adam Harouak, Jana Lenz, Aurelia Sahl, Maximilian Grohmann, Elisa Briem, Murray B Reed, Rodrig Marculescu, Lukas Nics, Sazan Rasul, Dan Rujescu, Marcus Hacker, Jens C Pruessner, Rupert Lanzenberger, Andreas Hahn

## Abstract

Psychosocial stress engages coordinated physiological and neural responses that enable adaptation to environmental challenges. Maladaptive stress responses and reduced resilience are major risk factors for psychiatric and neurodegenerative disorders. As the brain’s metabolic response to stress remains largely unexplored, we used simultaneous [^18^F]FDG PET/MRI during performance of the Montreal Imaging Stress Task to assess cerebral glucose metabolism, BOLD activation and functional connectivity. Multimodal data analysis identified the posterior cingulate cortex (PCC) as hub of stress processing beyond cognitive input, with increased glucose metabolism and attenuated BOLD deactivations. This was accompanied by reduced PCC connectivity within the default mode network and increased influence onto frontoparietal and dorsal attention networks. Endocrine stress responsivity was further reflected in lower resilience sub scores, failure to downregulate anterior cingulate cortex (ACC) metabolism during stress, and an inverse relationship between ACC glucose metabolism and anterior insula functional connectivity. These results indicate that acute psychosocial stress does not merely increase overall energy demand but reconfigures the relationship between regional metabolism and large-scale network organization, which further differs with individual endocrine stress responsivity. Our findings unify the roles of stress and the DMN in brain disorders, providing a multimodal framework for assessment in patient populations.

## Introduction

The response to stress is fundamentally adaptive, enabling maintained homeostasis when confronted with physical or psychological challenges.^1^ In humans the stress response is a coordinated activation of neuroendocrine, autonomic, and behavioural systems allowing rapid mobilization of metabolic resources.^1,2^ While acute stress responses are essential for immediate survival and considered adaptive, prolonged or dysregulated stress signalling has detrimental effects on body metabolism, brain function, and mental health. Here, substantial variability exists in individual stress responsivity, causing differences in the susceptibility for mental disease and resilience to psychosocial stressors.^2,3^ Maladaptive stress responsivity and impaired resilience play a critical role in the aetiology of psychiatric and neurodegenerative disorders, including major depressive disorder,^4,5^ schizophrenia,^6,7^ and Alzheimer’s disease, where psychosocial stress and neuroendocrine dysregulation have been implicated in disease pathogenesis and progression.^8^

A central component of the stress response is the hypothalamic-pituitary-adrenal (HPA) axis, coordinating endocrine signalling during stress.^1^ Perceived threat leads to the release of corticotropin-releasing hormone (CRH) from the hypothalamus, eliciting secretion of adrenocorticotropic hormone (ACTH) from the anterior pituitary. Subsequently, ACTH stimulates the adrenal cortex to secrete glucocorticoids, with cortisol being the principal stress hormone in humans.^1,3^ Cortisol acts on multiple peripheral and central targets to mobilize energy substrates, regulate metabolism, and modulate neural activity.^2,3,9^ Additionally, cortisol exerts negative feedback on the HPA axis via glucocorticoid receptors in brain regions including the prefrontal cortex (PFC), thereby limiting further hormone release.^10,11^

Consistent with its neuromodulatory effects, cortisol influences PFC function in part through glutamatergic signalling,^12^ which is closely linked to cellular energy metabolism.^13^ Synaptic release of glutamate increases glucose uptake in both neurons^14^ and astrocytes^15^ to support neurotransmitter recycling and restoration of ionic gradients.^13–15^ Accordingly, acute stress in rodents is associated with increased brain glucose metabolism, as measured using [^18^F]Fluorodeoxyglucose ([¹⁸F]FDG) positron emission tomography (PET), alongside elevated hexokinase activity in PFC synaptosomes.^16^ In non-human primates, different stress paradigms consistently increased glucose metabolism in the subgenual anterior cingulate cortex (ACC), with metabolic demands being positively associated with individual plasma cortisol levels.^17^

Despite the evidence from animal studies, characterization of stress-induced metabolic responses in the human brain are underexplored^18–20^, as neuroimaging of stress is mostly conducted using blood-oxygen-level-dependent (BOLD) functional magnetic resonance imaging (fMRI).^21^ Along with increased cortisol levels,^22^ these studies have shown that psychosocial stress robustly alters activation across cortical and limbic networks involved in cognitive control, emotional processing and threat assessment, including the ACC, pre- and orbitofrontal cortex, anterior insula, hippocampus and amygdala.^21–23^ These regions are implicated in emotional salience, evaluative processes and top-down regulation of the stress-response.^21,24^ Several of these regions, particularly medial prefrontal and cingulate areas, are integral parts of the default mode network (DMN), which is involved in self-referential processing and internally directed cognition.^25^ Increasing evidence suggests that the DMN is not simply suppressed during stress exposure but may actively contribute to stress-related processing, for instance through continued self-referential appraisal of the social-evaluative threat and rumination about one’s own performance.^26^ Notably, paradigms that require internally focused attention have repeatedly demonstrated a dissociation between glucose metabolism and BOLD-derived activity in the DMN,^27–30^ emphasizing the complementary information of the distinct imaging metrics, with the hemodynamic response of the BOLD signal and cellular energy demand, respectively.^31–34^ Altogether, these dissociations indicate that hemodynamic measures do not fully reflect the underlying metabolic demands,^27,28^ particularly that of stress-related glutamatergic activity.^35^ Thus, it is currently unknown whether a comparable divergence also emerges under acute psychosocial stress. Functional PET (fPET) imaging offers a direct approach to address this gap by resolving metabolic changes of multiple task conditions within a single scan, comparable to the acquisition of BOLD fMRI.^36,37^

The present study therefore aims to investigate the metabolic demands of acute psychosocial stress in humans, as well as corresponding BOLD signal changes and functional connectivity using simultaneous [^18^F]FDG PET/MRI. This multimodal imaging approach allowed us to investigate a potential mismatch between neurometabolic and neurovascular coupling mechanisms,^27^ as well as the influence of large-scale brain networks on glucose metabolism.^28^ By directly contrasting stress responders with non-responders based on the endocrine effects, we further assessed the neurobiological mechanisms of individual stress responsivity. For endocrine-related effects, the a priori hypothesis focused on the ACC, a structure particularly implicated in stress processing^17^ and involved in the emergence of psychiatric disorders.^19,21,38–40^

## Results

Sixty-six healthy volunteers underwent a protocol optimized to capture the stress response (figure 1a). This included a standardized light meal 5 hours before scanning to minimize variability of blood glucose, potentially influencing cortisol concentrations and [^18^F]FDG uptake^41,42^ as well as scheduling of measurements between 12:00 and 16:30 to reduce the influence of circadian fluctuations in cortisol levels. Venous cannulation for radiotracer application and blood sampling was performed 120 min before start of the PET/MRI session. The scan was preceded by a 60 min acclimatization period in the gantry, during which only MRI data were acquired while low-level visual stimuli were presented to ensure stress hormone levels had returned to baseline before task performance. Brain imaging comprised simultaneous acquisition of [^18^F]FDG fPET^36^ and BOLD fMRI while participants performed the Montreal Imaging Stress Task (MIST).^18,22^ In this paradigm participants solved simple mathematical calculations, first without (control condition) and then with time pressure and negative feedback (stress condition, figure 1b). Quantification of the cerebral metabolic rate of glucose (CMRGlu) was realized with a minimal-invasive protocol^43,44^ and venous blood samples were drawn for ACTH and cortisol levels. Different BOLD acquisitions were used as additional proxy of neuronal activation and for computation of stress-induced functional connectivity, respectively.

**Figure 1:**
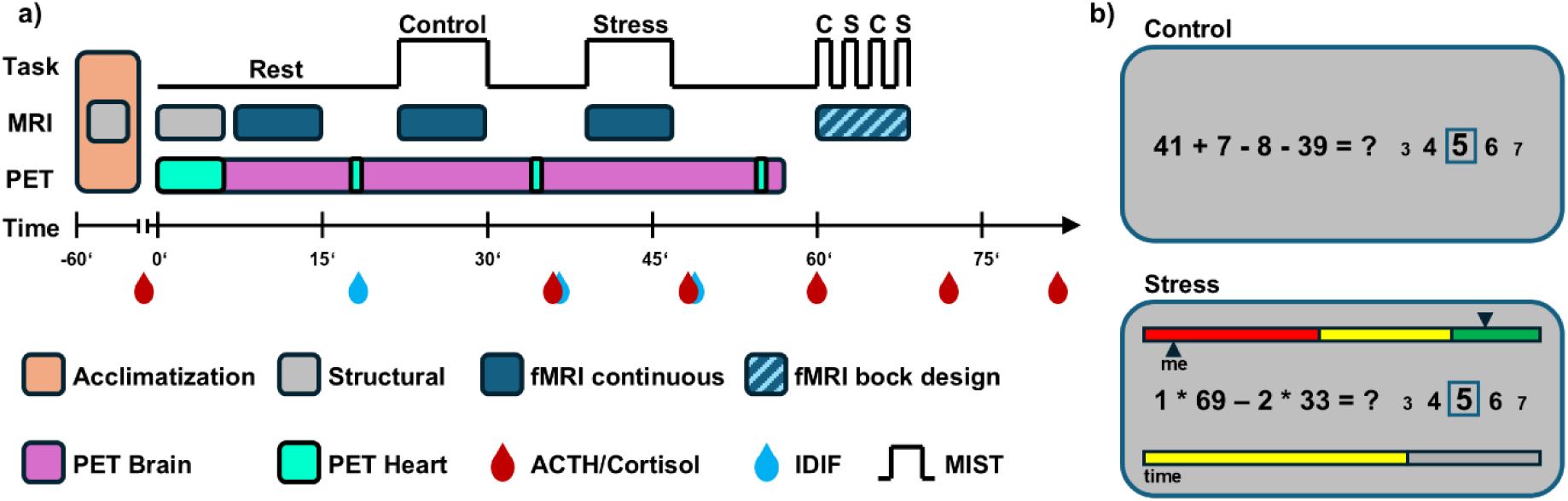
Study design. **(a)** Graphical overview of the simultaneous [¹⁸F]FDG PET/MRI protocol. Participants first underwent a 60-minute acclimatization period inside the scanner, during which low-level visual stimulation was presented and a structural MRI of the brain was acquired. The combined PET/MRI measurement then commenced with a bolus-plus-constant infusion of [¹⁸F]FDG (56 min). In the first six minutes, a structural T1-weighted MRI of the heart was additionally acquired for later use in image-derived input function (IDIF) calculation, with the scanner bed returning automatically to the heart at 17.5, 34 and 54.5 min. After an initial resting state period (8 min), participants then completed one control block and one psychosocial stress block of the Montreal Imaging Stress Task (MIST, 8 min each). The order of conditions was fixed, with the control condition always preceding the stress condition, as the endocrine response to a psychosocial stress induction cannot be reversed within a session. The herein acquired fMRI data was used for functional connectivity facilitated by the continuous task performance. After the PET acquisition, four additional MRI-only blocks were acquired in a conventional block design, which was used for estimation of neuronal activation (two control (C, 1 min), two stress (S, 2 min)). Throughout the scanning session, six venous blood samples were collected for quantification of ACTH and cortisol, and three additional samples were drawn to supplement the IDIF. **(b)** Example interface of the MIST as presented to participants. In the control condition (top), arithmetic problems were presented without a time limit and without performance feedback. In the stress condition (bottom), participants completed arithmetic problems under time pressure, indicated by a countdown bar, combined with manipulated performance comparison to an average control group and written feedback of the participant’s suboptimal performance designed to induce psychosocial stress.

### Endocrine, autonomic and behavioural stress reactions

To establish that the MIST elicited a physiological stress response, and to define individual stress responsivity for the imaging analyses, endocrine, autonomic and subjective markers were obtained throughout the scan session (figure 1a). ACTH was used as the primary marker of the endocrine stress response since the hormone is released from the anterior pituitary under direct control of hypothalamic CRH. Thus it provides a more direct correlate to brain activity and imaging parameters than cortisol, which is separated from the central drive of the HPA axis by a further adrenal step and by a considerably longer delay.^45^ Gaussian mixture modelling (GMM), a model-based clustering approach describing the distribution of ACTH response as a mixture of two latent Gaussian components,^46^ was used to differentiate between responders and non-responders. This identified 62.1% of participants as stress responders at a threshold of 25.8% increase in ACTH levels compared to pre-stress levels (supplementary figure 1a). This led to responders showing significantly higher stress-induced increases in ACTH (by definition of the GMM) and cortisol compared to non-responders for all time points post-stress (figure 2a-b, t= 2.8 to 5.7, p<0.05 to p<0.0001). Similarly, pulse recordings showed a significantly higher increase during stress for responders than non-responders (figure 2c, t=3.4, p<0.01). On a behavioral level, the subjective stress ratings right after the control and stress conditions increased for both responders and non-responders (figure 2d, both t>6.9, p<0.0001). The absence of a group difference in subjective ratings is consistent with the well-documented dissociation between subjective and physiological indices of the acute stress response.^47^ Still, responders showed lower resilience to stress at trend level for the full Connor-Davidson Resilience Scale (t=-1.8, p=0.08) and specifically for items of coping (mean±sd responder=14.8±2.6 vs. non-responder=16.1±2.0, t=-2.1, p<0.05) and meaningfulness (8.1±2.5 vs. 9.4±2.8, t=-2.0, p=0.05).

**Figure 2:**
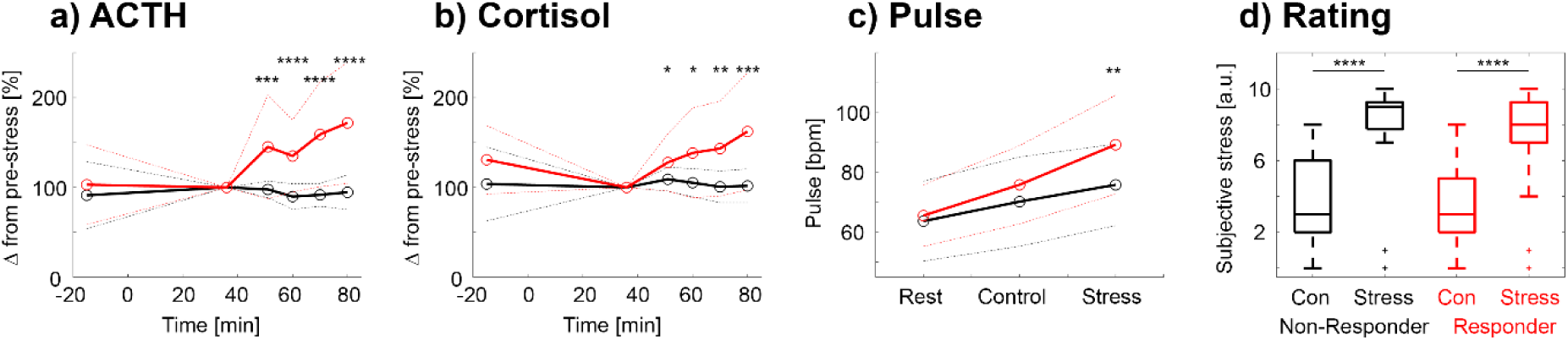
Endocrine, autonomic and subjective stress response. **a)** ACTH profiles in stress responders and non-responders. Group separation was obtained from percentage change in ACTH levels with Gaussian mixture modelling, yielding 62.1% of responders (n = 41 responders and n = 25 non-responders). Significant group differences were observed for all time points after stress exposure. b) Corresponding cortisol profiles showed similar characteristics, but less pronounced group separation. c) Heart rate increased more strongly during stress for responders than non-responders. d) Subjective stress ratings showed pronounced increases from control to stress conditions, but no difference between responders and non-responders. *p_Bonf_ < 0.05, **p_Bonf_ < 0.01, ***p_Bonf_ < 0.001, ****p_Bonf_ < 0.0001. red: stress responders, black: stress non-responders.

As cortisol represents a commonly used stress marker, results were replicated using an established cortisol response threshold of a 15% increase^48^ (supplementary figure 1b). Notably, this fixed, literature-based threshold for cortisol of 15% change is not directly comparable to the above data-driven cut-off for ACTH. Using cortisol data, 63.6% of participants were identified as stress responders, showing a high concordance with those identified using the ACTH based classification (χ^2^=17.4, p<0.0001). Moreover, separation between responders and non-responders was similarly effective for both the ACTH and cortisol threshold (both t>5.9, p<0.0001) and a moderate correlation between the two endocrine parameters was observed (supplementary figure 1c, rho=0.58, p<0.0001).

Taken together, the MIST elicited a robust endocrine and autonomic stress response with pronounced interindividual variability, which allowed a data-driven separation into responders and non-responders that was used for subsequent imaging analyses.

### Psychosocial stress distinctly increases DMN glucose metabolism and BOLD signal

Having established the physiological stress response, we next aimed to identify the corresponding metabolic and hemodynamic changes involved in stress processing. CMRGlu and BOLD signal changes were therefore quantified for the control and the stress condition of the MIST. During the control condition, activation in terms of CMRGlu and BOLD signal changes were observed predominantly in the visual, frontoparietal and dorsal attention networks (p<0.05 FWE corrected, supplementary figures S2 and S3). Similar to previous work,^49^ the two imaging modalities showed moderate spatial overlap of activations (Dice coefficient = 0.46). However, the PCC exhibited discordant changes, characterized by a decreased BOLD response, without corresponding differences in CMRGlu (supplementary figures S2 and S3).

Adding psychosocial stress to the task resulted in marked differences between the two modalities, with the Dice coefficient decreasing to 0.17 (figure 3a,d). Stress-specific increases in glucose metabolism were observed predominantly within the DMN, including subcortical representation in the thalamus,^50^ visual and frontoparietal networks (main effect of condition, p<0.05 FWE corrected). Of these stress-induced increases, only 19.8% of voxels overlapped with activations observed during the control condition and these were almost exclusively located in the visual cortex (supplementary figure S2), whereas the remaining regions were recruited *de novo* during stress (figure 3a-c). This stress-induced involvement of new areas contrasts with other tasks,^49,51^ where an increase in cognitive load results in a corresponding upregulation of CMRGlu, mostly in the same areas (i.e., 60.4% of areas already activated in the control condition (supplementary figure S4)). Findings on stress-induced metabolic demands remained virtually identical when using percent signal change^52^ instead of CMRGlu as an outcome parameter, indicating that these were genuinely driven by the stimulation, not by changes in baseline glucose metabolism (supplementary figure S5).

**Figure 3:**
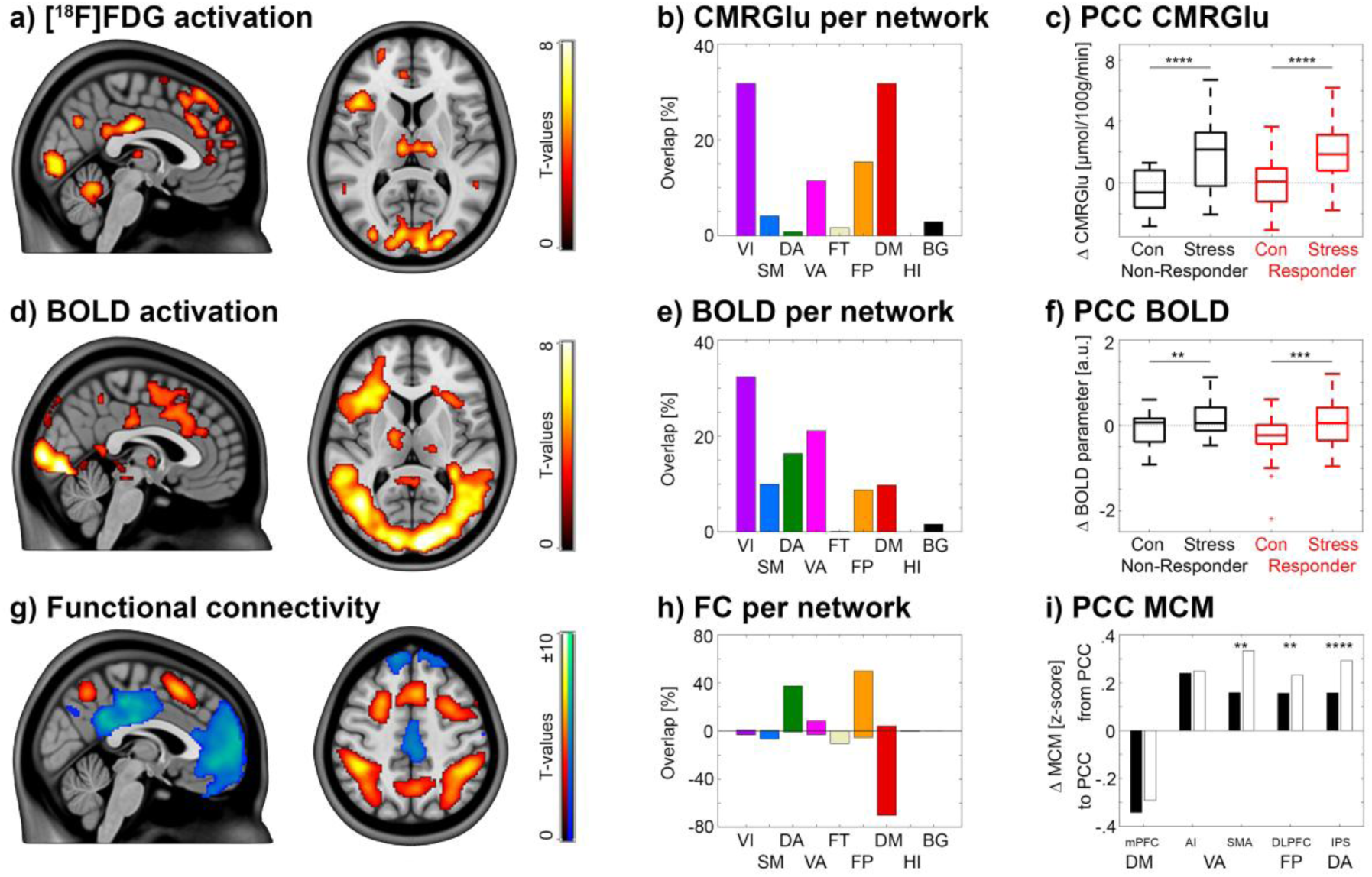
Stress-specific brain response. Images show the direct contrast of stress – control conditions (Δ). a-c) Glucose metabolism (CMRGlu) increased predominantly in the default mode and visual networks, particularly in the posterior cingulate cortex (PCC). These regions were not activated for the underlying cognitive task (supplementary figure S2) and thus newly recruited during stress. d-f) BOLD signal increases were observed in the visual and attention networks, as well as the PCC. In contrast to CMRGlu, BOLD deactivations of the PCC during the control condition (supplementary figure S2) were attenuated by stress. g-h) Proceeding from CMRGlu changes in the PCC as seed region, functional connectivity decreased within the DMN but increased for frontoparietal and dorsal attention networks. i) Directionality assessed with metabolic connectivity mapping (MCM) indicated that the PCC receives input from the mPFC, but exerts its influence on task-positive networks. This influence was higher during the stress condition for regions of the dorsal attention and frontoparietal networks. Results are corrected for multiple comparison (p<0.05 FWE for a, d, g; **p_Bonf_ < 0.01, ****p_Bonf_ < 0.0001 for c, f, i). red: stress responders, black: stress non-responders. i) black: control, white: stress.

Increases in the BOLD signal during stress were observed mostly among visual, dorsal and ventral attention networks (p<0.05 FWE corrected, figure 3d-f). In total, 40.5% of stress-specific activations overlapped with those of the control condition and were mostly distributed among these three networks (supplementary figure S3). Similar to CMRGlu, though less pronounced, the BOLD signal increased for stress vs. control in the DMN, specifically the PCC, which was the only network without observed BOLD activation during the control condition (figure 3d-f, supplementary figure S3). However, considering the distinct signal changes during the control condition (negative BOLD, unchanged CMRGlu), stress resulted in a return to baseline for the BOLD signal but a net positive CMRGlu response (figure 3d-f). Thus, stress-induced increases in metabolic demands of the PCC are also reflected in an increased hemodynamic response, independent of the initial condition of the underlying cognitive task.

### Stress shifts PCC connectivity from the DMN towards task-positive networks

Identifying the distinct role of the PCC in stress processing, we examined the corresponding effects with respect to its role in whole-brain network organization. Here, voxel-wise functional connectivity (FC) analysis was performed with the PCC as seed region. The continuous task performance during BOLD acquisition enabled us to investigate stress-induced changes in functional connectivity. Directionality of the resulting connections was subsequently inferred with metabolic connectivity mapping (MCM).^49,53^ This combination of FC and CMRGlu exploits the predominantly postsynaptic origin of energy consumption to identify the target region of a connection. Compared to the control condition, PCC connectivity within the DMN decreased, while FC with areas of the frontoparietal and dorsal attention networks increased (p<0.05 FWE corrected, figure 3g-h). MCM showed that the PCC receives input from the mPFC but exerts influence on task-positive networks. This influence significantly increased during the stress condition for the regions of the frontoparietal and dorsal attention networks (figure 3i, all p<0.05 corrected). Metabolic recruitment of the PCC was thus accompanied by a reorganization of its large-scale connectivity, with reduced coupling within the DMN and an increased influence on task-positive networks.

### ACC metabolism reflects individual stress response

Considering the established involvement of the ACC in stress processing,^17,19,24,40^ we specifically tested the *a priori* hypothesis that metabolic demands in this region vary with the individual stress response. While non-responders showed decreased CMRGlu during stress vs. control, responders showed increased metabolism, both in comparison to the control condition and to non-responders during stress (interaction effect, p<0.05 FWE corrected for the ACC, figure 4a-b). This finding was also replicated with the established cortisol-based threshold^48^ to separate stress responders from non-responders (supplementary figure S6). As expected from the above analysis, individual changes in ACTH levels were associated with stress-induced ACC CMRGlu values (rho=0.43, p<0.001). Still, the relationship was specific to the ACC as no comparable effect was observed for the PCC CMRGlu alone (rho=-0.14, p=0.3) and it remained stable even when adjusting for PCC metabolism (rho=0.43, p<0.001).

**Figure 4:**
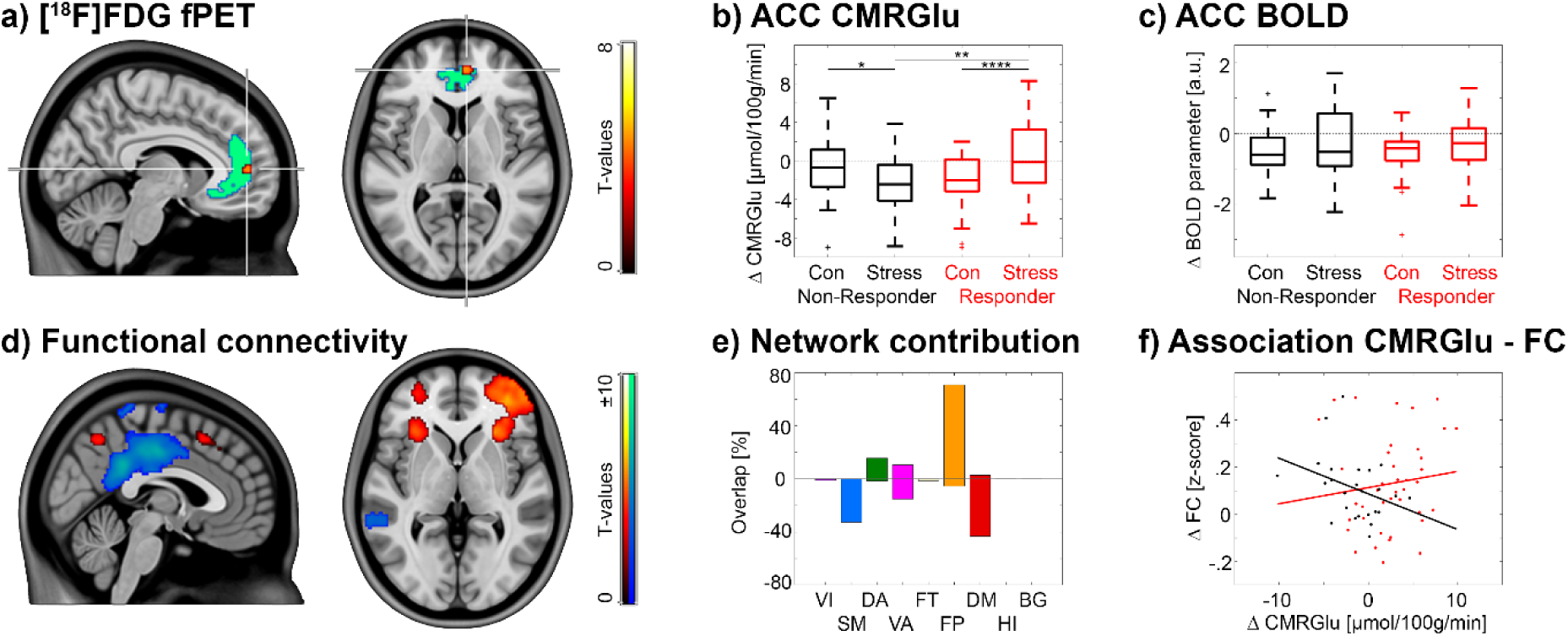
Differences between stress responders and non-responders. **a-b)** While glucose metabolism (CMRGlu) decreased during the stress condition for non-responders, stress-responders failed to downregulate CMRGlu in the anterior cingulate cortex (ACC, p<0.05 FWE corrected for the ACC region, shown in green). c) In contrast, no significant differences between groups or conditions were observed for the BOLD signal. d-e) Using the ACC as seed region, functional connectivity (FC) decreased within the DMN, but increased for the frontoparietal network and the anterior insula (p<0.05 FWE corrected). f) Stress responders and non-responders showed an inverse relationship between ACC CMRGlu and anterior insula FC (rho = 0.19 vs. −0.40, p<0.05). red: stress responders, black: stress non-responders.

For neuronal activation assessed with BOLD fMRI, no interaction effects were observed. Also, *post-hoc* exploratory analysis based on areas with CMRGlu effects, showed no significant changes in BOLD contrasts in the ACC between responder and non-responder neither for stress nor the control conditions (figure 4c).

Similar to the PCC, functional connectivity of the ACC was decreased within the DMN (p<0.05 FWE corrected, figure 4d) but predominantly increased within areas of the frontoparietal network (figure 4e). Notably, the anterior insula exhibited stress-specific increases in FC for both the general stress reaction (PCC seed, figure 3g) and the endocrine-driven stress response (ACC seed, figure 4d). Further evaluating the association between ACC CMRGlu and anterior insula FC showed an inverse relationship between responders and non-responders (rho=0.19 vs. −0.40, p<0.05, figure 4f). Individual endocrine stress responsivity was therefore mirrored in the metabolic regulation of the ACC and in its relation to salience network connectivity, in the absence of a corresponding effect in the BOLD signal.

## Discussion

Individual differences in stress vulnerability are increasingly understood to arise from distributed patterns of brain network organization.^54^ Consistent with this systems level perspective, psychosocial stress elicited coordinated neuroendocrine, metabolic, functional and network-specific responses that support behavioural adaptations to environmental challenges^1,2,21,24^ and extend beyond those previously described in studies using BOLD fMRI alone.^19,21,23,55^ Specifically, stress increased glucose metabolism within core regions of the DMN, while deactivation in the BOLD signal was attenuated. This was accompanied by decreased PCC input from the mPFC and increased output to task-positive networks. Moreover, individuals exhibiting an endocrine stress response scored at trend level lower on resilience, failed to downregulate ACC metabolism, and displayed an inverse relationship between ACC glucose metabolism and anterior insular functional connectivity.

The increased glucose metabolism in the DMN during psychosocial stress extends previous [^18^F]FDG PET work in rodents and non-human primates to humans.^16,17^ Stress paradigms in animals consistently showed increased cerebral glucose utilization within prefrontal regions implicated in stress regulation.^12,16,17,35^ In contrast, previous human fPET studies investigating cognitive performance (without explicit stress) reported upregulation of glucose metabolism as a function of cognitive load primarily in regions directly recruited by the task itself.^28^ Thus, our findings suggest that psychosocial stress additionally engages glucose metabolism in the DMN, which remains metabolically unchanged during cognitively demanding tasks alone,^28,56^ indicating sustained DMN engagement during social-evaluative threat.^57^ This observation contrasts with the classical view of the DMN as a task-negative network whose activity is suppressed during goal-directed cognition.^56,58,59^ One interpretation is that psychosocial stress recruits DMN functions including self-referential processing, rumination, and internal mentation during social evaluation.^25,26,56,58–60^ This is consistent with prior fMRI studies reporting altered DMN activity during stress and in disorders characterized by increased self-referential processing, including depression and anxiety.^26,60,61^

Interestingly, both PCC glucose metabolism and BOLD signal increased during stress, although from different levels as elicited by the control condition. The observed mismatch between neurometabolic (unchanged CMRGlu) and neurovascular processes (decreased BOLD contrast) of the underlying mathematical task alone matches previous reports during working memory.^27,28,30^ This has been interpreted as potential GABAergic inhibition, which implies high metabolic costs but decreases in the BOLD signal,^62–64^ emphasizing that glucose metabolism and BOLD signals capture complementary aspects of neuronal processing rather than representing redundant measures of brain activity.^27,30^ We further speculate that the subsequent increase in both signals is related to stress-induced glutamatergic neurotransmission, in line with preclinical work.^12,65^ This yields concurrent increases in local energetic demand and the BOLD signal ^16,66^, while inhibitory processes contribute to the observed dissociation between glucose metabolism and the BOLD signal.^27^

The stress-specific engagement of the PCC co-occurred with decreased within-network functional connectivity from the mPFC and increased connectivity towards task-positive networks involved in cognitive control and attention,^28,59,67^ with the directionality of these connections inferred from MCM. Thus, the overall change in network organization suggests greater functional integration^68^ and a shift from internally to externally oriented processing.^28,69^ This is in line with the interpretation that psychosocial stress elicits dynamic network reconfiguration supporting adaptation to evaluative threat,^1,2,5^ and with prior fMRI reports of altered coupling between the DMN and executive networks in states of stress and anxiety.^26,60,61^ Moreover, the combination of increased DMN metabolism and reduced within-network connectivity resembles alterations reported in psychiatric disorders associated with cognitive deficits, childhood trauma, and psychotic symptoms.^60,70,71^ Although the present study examined healthy participants, repeated recruitment of these metabolic and network-level adaptations may contribute to the network dysfunction observed in stress-related psychiatric disorders.^60,70^

Interindividual differences in endocrine stress responsiveness were accompanied by distinct metabolic and network-level brain responses, located in the anterior-medial part of the DMN. In individuals exhibiting a robust endocrine response to psychosocial stress, ACC glucose metabolism was elevated compared to the control condition, whereas non-responders showed a relative downregulation of ACC CMRGlu. These differences in metabolic demands may reflect the individual variability in regulatory processes during psychosocial stress, with the downregulation of ACC metabolism potentially indicating the successful engagement of inhibitory processes or control-related circuits.^72^ This interpretation was not directly tested by the present work and would require targeted assessment of inhibitory neurotransmission. Still, it aligns with previous studies implicating ACC function in stress sensitivity and affective disorders, particularly major depressive disorder (MDD) and anxiety disorders. Patients suffering from MDD show a blunted endocrine stress response, decreased ACC metabolism at rest,^39,73^ and decreased GABA levels in the ACC, indicating altered inhibitory processing.^72^ As the former two MDD findings are similar to healthy non-responders during stress, the observed group pattern is compatible with the possibility of an inverted U-shaped relationship between ACC metabolic demands and stress responsiveness. This hypothesis should be tested in future studies, directly comparing patients with healthy individuals in the same setting.

Interestingly, ACC metabolism was inversely associated with functional connectivity of the anterior insula between stress responders and non-responders. The anterior insula constitutes a key cortical node of the salience network, which is thought to detect behaviourally relevant stimuli and coordinate dynamic interactions between internally and externally oriented brain networks.^74–76^ The difference in the association between ACC CMRGlu and anterior insula FC potentially reflects variability in how salient social-evaluative information is processed and integrated during acute stress.^75,76^ Additionally, this interaction of the DMN and salience network further characterize higher-order network level organisation, potentially explaining stress-related alterations in interoception and emotional awareness.^77–79^ Considering that stress responders also showed a trend towards lower resilience on a behavioural level, our findings are in line with previous work suggesting that resilience modulates physiological and neural responses to stress.^80,81^ Together, these observations suggest that individual variability in stress responsiveness is associated with distinct alterations in endocrine signalling, glucose metabolism and functional connectivity that may contribute to vulnerability to stress-related psychopathology.

While simultaneous [^18^F]FDG PET/MRI is an effective design, which allowed a multimodal characterization of the response to acute psychosocial stress in humans, several limitations should be considered. Individual differences in stress perception may have contributed to the variability of the endocrine response. As a strength, this variability enabled the direct comparison of responders and non-responders. We cannot exclude, however, that individual non-responders recognized the manipulated nature of the task, as this was not assessed systematically. Despite several precautions before the scan (e.g., standardized meal, acclimatization, no arterial sampling), the experimental setting itself constitutes a limitation.

Participants underwent stress induction within the confines of a PET/MRI scanner, characterized by an unfamiliar environment, the requirement to remain still, and the presence of venous lines for radiotracer administration and blood sampling. This may differ from the psychosocial stressors typically encountered in everyday life, and the extent to which our findings extrapolate to daily psychosocial stress remains to be determined. Furthermore, although the present study identifies distinct alterations in cerebral glucose metabolism and functional connectivity in healthy participants during acute psychosocial stress, the cross-sectional design in healthy volunteers does not allow conclusions about the development of stress-related psychiatric disorders. The distinct metabolic and hemodynamic profiles of responders and non-responders should therefore be examined in clinical populations, in order to determine whether they map onto a consistent link with stress-related psychopathology. As the endocrine response to a psychosocial stress induction cannot plausibly be reversed within a session, the study was not designed as a counterbalanced cross-over between control and stress conditions. Order effects such as time on task or global drifts in tracer uptake can therefore not be ruled out in full. Such effects are however unlikely given the regionally specific of stress (instead of a global increase in task-specific CMRGlu). Also, the circumstance that findings were virtually unchanged when using percent signal change instead of CMRGlu argues against a purely global metabolic drift.

In conclusion, acute psychosocial stress is associated with coordinated changes in regional glucose metabolism, neuronal activation, and large-scale functional network organization, with the DMN taking a central role. This multimodal framework opens several avenues for future investigations. The distinct neurometabolic and neurovascular responses in the PCC during cognitive and stress processing warrant closer investigation of the underlying neurotransmitter systems. Extending this protocol to patient cohorts, the altered regulation of ACC metabolism may offer a target for assessing individual stress responsiveness and therapeutic response in MDD and other internalizing disorders such as anxiety disorders.

## Methods

### Experimental design

All participants completed a simultaneous PET/MRI measurement with the radiotracer [^18^F]FDG. The scan session was optimized to robustly capture the stress response. Participants were instructed to maintain a regular sleep-wake cycle at least three days prior to each appointment and refrain from sports on the day of the measurement. To minimize potential stress effects related to prolonged fasting, participants consumed a standardized light meal (salad or yogurt) 5 hours before stress exposure. In addition, they fasted three hours before this meal and between the meal and the PET/MRI scan, except for unsweetened water. Imaging sessions started between 12:00 and 16:30 to minimize circadian rhythm influence on cortisol. Moreover, venous cannulation for radiotracer application and blood sampling was done 120 min before start of the PET/MRI session and the scan was preceded by a 60 min acclimatization period in the gantry to ensure stress hormones levels return to baseline (MRI acquisition only, low-level visual input).^37,82^ After the acclimatization period, subjects were asked to visit the bathroom to avoid discomfort and movement in the subsequent scan. At this time point, blood glucose levels were measured for quantification of CMRGlu and baseline cortisol and ACTH samples were obtained.

PET/MRI acquisition was carried out with a shuttle protocol (70 min in total, figure 1a), which enables capturing a cardiac image-derived input function for non-invasive quantification of CMRGlu as well as fPET brain imaging.^44^ Accordingly, the first six minutes after radiotracer application, as well as three 1.5 minute periods at 17.5, 34 and 54.5 min were acquired at the cardiac position, and the remainder of the scan at the brain position. During the scan, participants completed the MIST, with control and stress conditions starting 23 and 40 min after radiotracer application, respectively (8 min each). Task performance was carried out continuously (i.e., without rest-task transitions) and BOLD fMRI was acquired simultaneously to facilitate computation of task-specific functional connectivity.^49^ After the fPET acquisition, a final BOLD fMRI sequence was obtained, with MIST performance in a block design similar to previous work^22,83^ and used as a proxy of neuronal activation (2x 60 s control blocks, 2x 120 s stress blocks, alternating order, 30 s baseline before, between and after task blocks, 8.5 min in total). Thus, in a strict sense only the continuous BOLD acquisition for functional connectivity was obtained simultaneously with fPET, whereas the block-design BOLD acquisition used to estimate neuronal activation followed the fPET measurement.

Blood samples were collected during rest periods to assess plasma cortisol and ACTH levels (pre-stress: −25, 36 min; post-stress: 50.5, 60, 70, 80 min relative to scan start) and to supplement the IDIF (19.5, 36, 50.5 min). Heart rate was continuously monitored during baseline and task performance to assess autonomous stress response. After the control and the stress tasks, participants rated the subjective stress experience of each condition on a scale from 0 to 10.

### Participants

Seventy-two healthy participants were recruited and after inclusion invited for the scan session. In six cases scans could not be completed due to discomfort (3), image quality (2) or acute lack of radiotracer (1). Thus, 66 healthy participants (30 female; mean age±SD = 24.1±4.7 years) were included in this analysis. They underwent a routine medical examination during a screening visit including electrocardiography, standardized blood test, neurological and physical examination. Furthermore, a urine drug test as well as a pregnancy test for female participants was performed at screening and on the day of the scan. Psychiatric disorders were excluded using the Structured Clinical Interview for DSM-V (SCID) by an experienced interviewer. Participants were eligible for inclusion if they were between 18 and 40 years of age, right-handed (to minimize potential hemispheric lateralization effects), and willing and able to provide written informed consent. Exclusion criteria comprised a current or past history of physical, neurological, or psychiatric disorders; current substance abuse or use of medication including antipsychotic, antidepressant or anxiolytic medication; pregnancy or breastfeeding; contraindications to MRI (e.g., metallic implants, including dental implants causing significant imaging artifacts); cumulative research-related exposure to ionizing radiation exceeding 30 mSv within the preceding 10 years; or inability to comply with the study protocol or follow instructions provided by the investigating team. At the screening visit participants also completed the Conor-Davidson Resilience Scale^84^ with permission by the original authors. After detailed explanation of the study protocol all participants gave written informed consent. All participants were insured and reimbursed for their participation. The study was approved by the ethics committee of the Medical University of Vienna (ethics number: 1642/2022) and all procedures were performed in accordance with the declaration of Helsinki. The study was registered to clinicaltrials.gov (NCT06243783).

### Montreal Imaging Stress Task (MIST)

The MIST is an adapted version of the Trier Social Stress Test optimized for imaging studies.^22^ The MIST consisted of a control and stress condition, which were both administered continuously for 8 minutes. During both conditions participants performed a mental arithmetic task consisting of up to four numbers (ranging from 0 to 99) and different mathematical operators (+, −, ×, ÷) (e.g., 12 × 6 - 10 × 7). The levels of task difficulty were slightly adapted from its original version.^22^ Level 1 comprised two numbers and only + or - operators. Level 2 included 3 numbers and optionally the × operator. Level 3 comprised 4 numbers. In level 4, the × operator was at least used once and in level 5 the ÷ operator was always included. The different levels of difficulty were presented in pseudo-randomized order to balance the appearance. All calculations were solved by an integer from 0 to 9, which participants were instructed to select using a button box with their right hand only (figure 1b). All calculations were tested for consistency and reaction times prior to inclusion in the task.

During the control condition, no stress-inducing elements were added. Participants were informed that their performance during this condition would not be assessed, no feedback about correctness was provided and no time limit was imposed. As a result, performance rates in the control condition were approximately 90%.

For the stress condition, a time limit, negative feedback and performance of an average control group were added. Participants were informed that the task was designed to investigate brain activation during mathematical calculations and that the average performance ranges between 80% and 90% correct answers and that a minimum accuracy of 80% was required. To enhance stress induction, this expectation was repeated immediately prior to the task. Unbeknownst to participants, the task was adaptively coded to maintain performance below this expected level. Specifically, the time limit of each level of difficulty was dynamically adjusted to 10% below the participant’s average response time of the control condition. This resulted in an overall success rate of approximately 40–50%. Additionally, the time limit of each level of difficulty was increased or decreased by 10% following three consecutive incorrect or correct responses, respectively. Stress was further augmented through the presentation of a performance bar comparing the performance of the participant and an average of an alleged control group, the latter exhibiting a success rate of 80–90%. Additionally, participants received negative feedback shown on the screen between trials, including reminders of their suboptimal and the required performance level as well as references to evaluation by investigators outside the scanner room. Following each condition, participants rated their perceived stress using a visual analogue scale. Participants were informed about the adaptive nature of the task or the stress manipulation only after study completion.

### PET/MRI acquisition

Imaging was performed at the Department of Radiology and Nuclear Medicine, Medical University of Vienna using a hybrid PET/MR system (Biograph mMR, Siemens Healthineers, Erlangen, Germany), equipped with a 12-channel head coil. The PET acquisition started 1 minute before radiotracer application to facilitate subsequent workflow and timings, but this initial time was subsequently discarded from the scan. The radiotracer [¹⁸F]FDG was administered using a bolus-plus-constant-infusion protocol, with 20% of the activity delivered as an initial bolus over 1 minute and the remaining activity infused continuously over 55 minutes (total of 5.1 MBq/kg body weight).^37^ For estimation of the image-derived input function, the bed position alternated between head and chest throughout the scan.^44^ At the chest position, a T1-weighted STARVIBE sequence (TE/TR = 1.44/3050 ms, flip angle = 5°, matrix size = 320 × 320, 208 slices, voxel size = 1.19 × 1.19 × 1.2 mm³, TA = 5:33 min) was obtained for cardiac imaging, together with a Dixon-based MR attenuation correction (MRAC) sequence using a CAIPIRINHA sampling pattern^85^ for attenuation correction of the chest PET data. A high-resolution structural MRI was acquired using a T1-weighted magnetization-prepared rapid gradient echo (MPRAGE) sequence (TE/TR = 4.21/2200 ms, voxel size = 1 × 1 × 1 mm³ + 0.1 mm gap, 240 × 256 × 160 slices). This was used to exclude gross anatomical abnormalities and for spatial normalization of PET data. BOLD fMRI data were acquired during continuous and blocked task performance using an echo-planar imaging (EPI) sequence (TE/TR = 30/2000 ms, voxel size = 2.5 × 2.5 × 2.5 mm³ with a 0.825 mm gap, 80 × 80 × 34 slices).

### Blood sampling and processing

Blood samples were drawn for determination of ACTH and cortisol plasma levels and to supplement the IDIF (see section experimental design for sample timings). The shorter plasma half-life of ACTH requires rapid subsequent processing, which was accounted for by immediate processing of every ACTH and cortisol sample at the internal laboratory of the University Hospital. For IDIF samples, whole-blood activity and (after centrifugation) plasma activity were measured in a gamma counter (Wizard^2^, 3”, PerkinElmer, Waltham, MA, USA), which was cross-calibrated with the PET/MRI system. An image-derived input function (IDIF) was obtained from PET images of the heart.^43,44^ Here, fixed-size regions of interest were manually placed in the ascending and descending aorta and the left ventricle. The IDIF peak was extracted from the descending aorta using the first 6 min of the scan in the cardiac position. The IDIF tail was given by the average values extracted from all three regions and the venous samples. The final input function was generated by scaling the whole-blood IDIF with the average plasma-to-whole blood ratio.^44^

### Quantification of glucose metabolism (CMRGlu)

Reconstruction and preprocessing of fPET brain images as well as quantification of CMRGlu was done as described previously.^49^ Image reconstruction was performed using an Ordinary Poisson Ordered Subset Expectation Maximization (OP-OSEM) algorithm (3 iterations, 21 subsets), resulting in images with a matrix size of 344 × 344 × 127 and a voxel size of 2.09 × 2.09 × 2.03 mm³. Dynamic data were reconstructed into 30-second frames. Standard corrections, including decay, scatter, and dead time, were applied, and attenuation correction was performed using a pseudo-CT approach derived from structural MRI.^86,87^ Motion correction and spatial preprocessing were performed using SPM12 (Statistical Parametric Mapping, Release 12, Wellcome Centre for Human Neuroimaging, London, UK, https://www.fil.ion.ucl.ac.uk/spm). Here, fPET images were realigned to the mean image (quality setting = 1), co-registered to the individual structural MRI, spatially normalized to MNI space using transformations derived from the anatomical scan and spatial smoothing with an 8 mm Gaussian kernel. To identify task-induced changes in glucose metabolism, fPET data were further analysed using the fPET Toolbox.^88^ A voxel-wise general linear model (GLM) was applied to separate task-related changes in glucose metabolism from baseline activity. Task regressors (baseline, control, stress, movement) were used to model dynamic changes in tracer uptake, with its slope reflecting task-specific alterations in glucose metabolism.^36^ The baseline regressor was defined as the average across all grey matter voxels, excluding those regions obtained in a meta-analysis of stress-specific neuronal activation.^24,37^ The two task regressors were constructed as a ramp function with a slope of 1 kBq/min during task performance and zero otherwise. The movement regressors were represented by the first principal components of the six movement regressors obtained during realignment. The net influx constant (K_i_) was estimated using the Gjedde-Patlak plot separately for each condition, assuming linearity after 20 minutes post-injection. K_i_ values were subsequently converted to CMRGlu with a lumped constant of 0.89.^36^

### Assessment of BOLD-derived neuronal activation

BOLD data preprocessing was done as described previously using SPM12.^49^ Data were corrected for slice timing (reference = middle slide) and head motion (quality setting = 1), spatially normalized to MNI space and smoothed with an 8 mm Gaussian kernel. Estimation of task-induced BOLD signal changes was done with the fMRI acquisition at the end of the scan, with performance of the MIST in a block design (figure 1). Here, a GLM was applied with two regressors modelling task effects (control and stress conditions), six for head motion and five for nuisance signals of the white matter and cerebrospinal fluid. Due to the longer task duration of the MIST,^22,83^ the high pass filter was set to 420 s.

### Computation of functional connectivity (FC) and metabolic connectivity mapping (MCM)

BOLD data preprocessing was completed in the same way as for neuronal activation until smoothing. Here, data from the 8 min continuous task performance of the control and stress conditions were used (figure 1). Then, motion censoring was applied using the DVARS approach^89^ as this provides a reasonable trade-off between signal denoising, data retention and computational complexity.^90^ Potentially confounding effects were removed by regression against realignment parameters as well as signals from white matter and cerebrospinal fluid, followed by bandpass filtering (0.01 < f < 0.15 Hz).^49,91^

Functional connectivity (FC) analyses were performed using a seed-based correlation approach, focusing on regions with group differences in CMRGlu (i.e., PCC and ACC, p<0.05 FWE corrected, see results). FC maps were generated by computing voxel-wise correlations of BOLD signal time courses between the seed region and all other brain voxels, followed by Fisher’s r-to-z transformation. This was done separately for control and stress conditions and the resulting FC maps were used for group level analyses.

Metabolic connectivity mapping (MCM)^53^ was computed on a region of interest basis as described previously.^28,49^ The main assumption is to infer directionality of functional connections as metabolic demands mainly arise postsynaptically,^33,34,92^ thereby identifying the target region of a connection. Regions were defined as exhibiting significant changes in CMRGlu (i.e., the PCC) and corresponding effects in FC (mPFC, anterior insula, DLPFC, IPS and SMA, see results). As the effect in the ACC was confined to a small area < 100 voxels, MCM was not computed with this target area to avoid spurious results due to limited spatial correlations. MCM employs several steps. First, FC is computed between regions A and B, yielding a voxel wise FC pattern in the putative target region B. This pattern is then spatially correlated with the corresponding voxel-wise pattern of CMRGlu in region B. Next, regions A and B are swapped and the calculation is repeated. The region with the significantly higher spatial correlation between FC and CRMGlu is then identified as the target region, enabling the calculation of directional connectivity.

### Statistical analysis

Statistics were computed using Matlab (R2018b, MathWorks, Natick, USA) and SPM12, where applicable. ACTH (and cortisol) levels were converted to percent change relative to pre-stress levels and Gaussian mixture modelling was used to separate responders from non-responders. For this model, subjects with extensive changes in ACTH levels (>142%) were not used in the model to avoid bias in the cut-off between groups. The value of 142% was defined from a conventional boxplot as the upper whisker, i.e., third quartile + 1.5 * interquartile range. ACTH and cortisol levels were compared between responders and non-responders using two-sample t-tests with Bonferroni correction for 4 post-stress time points. The relationship between responders and non-responders as defined by ACTH levels from Gaussian mixture modelling and the conventional 15% increase in cortisol levels^48^ was assessed by Chi-square test. Further, Pearson’s correlation was used to directly compare ACTH and cortisol levels. Pulse levels were compared between groups with two-sample t-test (Bonferroni corrected, 3 time points). Subjective stress ratings were compared between control and stress conditions using paired t-tests (Bonferroni corrected, 2 groups). Scores on the Connor-Davidson Resilience Scale were compared between groups using two-sample t-test.

Voxel-wise statistics in SPM12 were done in the same manner for CMRGlu, BOLD changes and FC with p<0.05 FWE corrected for multiple comparisons at cluster level following p<0.001 uncorrected voxel level. A repeated measures ANOVA was conducted with the between-subject factor group (levels: responder, non-responder), the within-subject factor condition (levels: control, stress) and subject as the repeated factor. Interaction effects and main effects of condition were assessed. One sample t-tests were used to assess the overall task effects as compared to rest. The Dice coefficient was used to compare the spatial overlap of activation maps between stress-induced changes in CMRGlu and the BOLD signal. Differences in MCM between conditions were assessed by paired t-test (Bonferroni corrected, 5 connections).

For the anterior cingulate cortex (ACC), the *a priori* hypothesis of differences between responders and non-responders was tested by small volume correction. Here, a similar repeated measures ANOVA model as above was used with the ACC region defined by the Mindboggle brain atlas^93^ and the statistical threshold was set to p<0.05 FWE corrected voxel level. For the association between ACC CMRGlu and anterior insula FC (see results), imaging metrics were extracted from the significant clusters, and Spearman’s correlation was used to avoid bias of potential outliers. Significant differences in correlations were assessed by z-transformation (R_1_’, R_2_’) and subsequent computation of a Z statistic as done previously,^94^ with n1 and n2 representing the number of subjects in each group (Equ. 1). The subtraction of three in the denominator reflects the variance of the Fisher z-transformed correlation coefficient, which is given by 1/(n - 3).

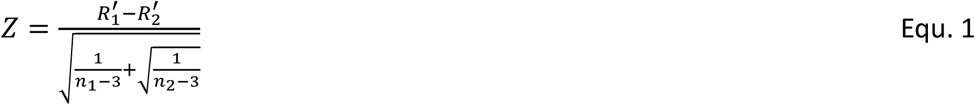

## Acknowledgments

We thank the graduated team members and the diploma students of the Neuroimaging Lab (NIL, head R. Lanzenberger) as well as the clinical colleagues from the Department of Psychiatry and Psychotherapy for clinical and/or administrative support, especially B. Eggerstorfer, M. Ponce de Leon and R. Röhheuser. We are particularly grateful to Y Qiu, T.M.C. Lee and R. Huang for providing the ALE meta-analysis maps of brain activation during acute stress. The scientific project was performed with the support of the Research Platform Medical Imaging of the Medical University of Vienna.

## Author contribution statement (CRediT)

**Gabriel Schlosser:** Investigation, Writing – original draft, Visualisation **Christian Milz:** Investigation, Data curation, Funding acquisition **Pia Falb:** Investigation, Data curation **Samantha Graf:** Investigation **Sarah M Tüchler:** Investigation, Data curation **Matej Murgaš**: Investigation, Data curation **Alexandra Mayerweg:** Investigation **Clemens Schmidt:** Investigation **Ivan Pörnbacher**: Investigation **Lukas Artmeier:** Investigation **Adam Harouak:** Investigation **Aurelia Sahl:** Project administration **Maximilian Grohmann**: Investigation **Murray B Reed:** Software, Validation, Data curation **Elisa Briem:** Investigation, Funding acquisition **Rodrig Marculescu:** Validation, Resources **Lukas Nics:** Supervision, Project administration **Sazan Rasul:** Supervision **Dan Rujescu:** Supervision, Resources **Marcus Hacker:** Supervision, Resources **Jens C Pruessner:** Conceptualization, Methodology **Rupert Lanzenberger:** Conceptualization, Supervision, Resources, Project administration, Funding acquisition **Andreas Hahn:** Conceptualization, Methodology, Software, Validation, Formal analysis, Visualization, Supervision, Funding acquisition, Writing – original draft. **All authors:** Writing - Review & Editing

## Ethical considerations

The study was approved by the Ethics Committee of the Medical University of Vienna (ethics number: 1642/2022) and all procedures were carried out in accordance with the Declaration of Helsinki. The study was pre-registered at clinicaltrials.gov with the number NCT06243783.

## Consent to participate

After detailed explanation of the study protocol, all subjects gave written informed consent. All subjects were insured and reimbursed for their participation.

## Consent for publication

Not applicable.

## Data availability

Raw data will not be publicly available due to reasons of data protection. Processed data and custom code can be obtained from the corresponding author with a data sharing agreement, approved by the departments of legal affairs and data clearing of the Medical University of Vienna.

## References

1. Ulrich-Lai, Y. M. & Herman, J. P. Neural Regulation of Endocrine and Autonomic Stress Responses. Nat. Rev. Neurosci. 10, 397–409 (2009).

2. McEwen, B. S. Physiology and neurobiology of stress and adaptation: central role of the brain. Physiol. Rev. 87, 873–904 (2007).

3. de Kloet, E. R., Joëls, M. & Holsboer, F. Stress and the brain: from adaptation to disease. Nat. Rev. Neurosci. 6, 463–475 (2005).

4. Krishnan, V. & Nestler, E. J. The molecular neurobiology of depression. Nature 455, 894–902 (2008).

5. Lupien, S. J., McEwen, B. S., Gunnar, M. R. & Heim, C. Effects of stress throughout the lifespan on the brain, behaviour and cognition. Nat. Rev. Neurosci. 10, 434–445 (2009).

6. Howes, O. D. & Murray, R. M. Schizophrenia: an integrated sociodevelopmental-cognitive model. Lancet 383, 1677–1687 (2014).

7. van Os, J., Kenis, G. & Rutten, B. P. F. The environment and schizophrenia. Nature 468, 203–212 (2010).

8. Milligan Armstrong, A., et al. Chronic stress and Alzheimer’s disease: the interplay between the hypothalamic-pituitary-adrenal axis, genetics and microglia. Biol. Rev. Camb. Philos. Soc. 96, 2209–2228 (2021).

9. Zhu, L.-J. et al. The Different Roles of Glucocorticoids in the Hippocampus and Hypothalamus in Chronic Stress-Induced HPA Axis Hyperactivity. PLoS ONE 9, e97689 (2014).

10. Arnsten, A. F. T. Stress signalling pathways that impair prefrontal cortex structure and function. Nat. Rev. Neurosci. 10, 410–422 (2009).

11. Herman, J. P. et al. Regulation of the Hypothalamic-Pituitary-Adrenocortical Stress Response. Compr. Physiol. 6, 603–621 (2016).

12. Musazzi, L. et al. Acute Stress Increases Depolarization-Evoked Glutamate Release in the Rat Prefrontal/Frontal Cortex: The Dampening Action of Antidepressants. PLoS ONE 5, e8566 (2010).

13. Attwell, D. et al. Glial and neuronal control of brain blood flow. Nature 468, 232–243 (2010).

14. Lundgaard, I. et al. Direct neuronal glucose uptake heralds activity-dependent increases in cerebral metabolism. Nat. Commun. 6, 6807 (2015).

15. Zimmer, E. R. et al. [18F]FDG PET signal is driven by astroglial glutamate transport. Nat. Neurosci. 20, 393–395 (2017).

16. Musazzi, L. et al. Acute Inescapable Stress Rapidly Increases Synaptic Energy Metabolism in Prefrontal Cortex and Alters Working Memory Performance. Cereb. Cortex 29, 4948–4957 (2019).

17. Jahn, A. L. et al. Subgenual prefrontal cortex activity predicts individual differences in hypothalamic-pituitary-adrenal activity across different contexts. Biol. Psychiatry 67, 175–181 (2010).

18. Pruessner, J. C., Champagne, F., Meaney, M. J. & Dagher, A. Dopamine release in response to a psychological stress in humans and its relationship to early life maternal care: a positron emission tomography study using [11C]raclopride. J. Neurosci. Off. J. Soc. Neurosci. 24, 2825–2831 (2004).

19. Pruessner, J. C. et al. Deactivation of the limbic system during acute psychosocial stress: evidence from positron emission tomography and functional magnetic resonance imaging studies. Biol. Psychiatry 63, 234–240 (2008).

20. Kern, S. et al. Glucose metabolic changes in the prefrontal cortex are associated with HPA axis response to a psychosocial stressor. Psychoneuroendocrinology 33, 517–529 (2008).

21. Dedovic, K., D’Aguiar, C. & Pruessner, J. C. What stress does to your brain: a review of neuroimaging studies. Can. J. Psychiatry Rev. Can. Psychiatr. 54, 6–15 (2009).

22. Dedovic, K. et al. The Montreal Imaging Stress Task: using functional imaging to investigate the effects of perceiving and processing psychosocial stress in the human brain. J. Psychiatry Neurosci. JPN 30, 319–325 (2005).

23. Kogler, L. et al. Psychosocial versus physiological stress - Meta-analyses on deactivations and activations of the neural correlates of stress reactions. NeuroImage 119, 235–251 (2015).

24. Qiu, Y. et al. Brain activation elicited by acute stress: An ALE meta-analysis. Neurosci. Biobehav. Rev. 132, 706–724 (2022).

25. Davey, C. G., Pujol, J. & Harrison, B. J. Mapping the self in the brain’s default mode network. NeuroImage 132, 390–397 (2016).

26. Hamilton, J. P., Farmer, M., Fogelman, P. & Gotlib, I. H. Depressive Rumination, the Default-Mode Network, and the Dark Matter of Clinical Neuroscience. Biol. Psychiatry 78, 224–230 (2015).

27. Stiernman, L. J. et al. Dissociations between glucose metabolism and blood oxygenation in the human default mode network revealed by simultaneous PET-fMRI. Proc. Natl. Acad. Sci. U. S. A. 118, e2021913118 (2021).

28. Godbersen, G. M. et al. Task-evoked metabolic demands of the posteromedial default mode network are shaped by dorsal attention and frontoparietal control networks. eLife 12, e84683 (2023).

29. Putkinen, V. et al. Cerebral glucose utilisation during musical emotions: A multimodal functional PET/MRI study. NeuroImage 338, 122035 (2026).

30. Hahn, A. et al. High-temporal resolution functional PET/MRI reveals coupling between human metabolic and hemodynamic brain response. Eur. J. Nucl. Med. Mol. Imaging 51, 1310–1322 (2024).

31. Logothetis, N. K. & Pfeuffer, J. On the nature of the BOLD fMRI contrast mechanism. Magn. Reson. Imaging 22, 1517–1531 (2004).

32. Heeger, D. J. & Ress, D. What does fMRI tell us about neuronal activity? Nat. Rev. Neurosci. 3, 142–151 (2002).

33. Mergenthaler, P., Lindauer, U., Dienel, G. A. & Meisel, A. Sugar for the brain: the role of glucose in physiological and pathological brain function. Trends Neurosci. 36, 587–597 (2013).

34. Harris, J. J., Jolivet, R. & Attwell, D. Synaptic energy use and supply. Neuron 75, 762–777 (2012).

35. Musazzi, L., Treccani, G. & Popoli, M. Functional and structural remodeling of glutamate synapses in prefrontal and frontal cortex induced by behavioral stress. Front. Psychiatry 6, 60 (2015).

36. Hahn, A. et al. Quantification of Task-Specific Glucose Metabolism with Constant Infusion of 18F-FDG. J. Nucl. Med. Off. Publ. Soc. Nucl. Med. 57, 1933–1940 (2016).

37. Rischka, L. et al. Reduced task durations in functional PET imaging with [18F]FDG approaching that of functional MRI. NeuroImage 181, 323–330 (2018).

38. Hamilton, J. P. et al. Functional neuroimaging of major depressive disorder: a meta-analysis and new integration of base line activation and neural response data. Am. J. Psychiatry 169, 693–703 (2012).

39. Drevets, W. C. et al. Subgenual prefrontal cortex abnormalities in mood disorders. Nature 386, 824–827 (1997).

40. Dolfen, N. et al. Stress Modulates the Balance between Hippocampal and Motor Networks during Motor Memory Processing. Cereb. Cortex 31, 1365–1382 (2021).

41. von Dawans, B., Zimmer, P. & Domes, G. Effects of glucose intake on stress reactivity in young, healthy men. Psychoneuroendocrinology 126, 105062 (2021).

42. Sarikaya, I., Sarikaya, A. & Sharma, P. Assessing the Effect of Various Blood Glucose Levels on 18F-FDG Activity in the Brain, Liver, and Blood Pool. J. Nucl. Med. Technol. 47, 313–318 (2019).

43. Reed, M. B. et al. Comparison of cardiac image-derived input functions for quantitative whole body [18F]FDG imaging with arterial blood sampling. Front. Physiol. 14, 1074052 (2023).

44. Reed, M. B. et al. Validation of cardiac image-derived input functions for functional PET quantification. Eur. J. Nucl. Med. Mol. Imaging (2024) doi:10.1007/s00259-024-06716-8.

45. Bae, Y. J. et al. Salivary cortisone, as a biomarker for psychosocial stress, is associated with state anxiety and heart rate. Psychoneuroendocrinology 101, 35–41 (2019).

46. General Introduction. in Finite Mixture Models 1–39 (John Wiley & Sons, Ltd, 2000). doi:10.1002/0471721182.ch1.

47. Campbell, J. & Ehlert, U. Acute psychosocial stress: does the emotional stress response correspond with physiological responses? Psychoneuroendocrinology 37, 1111–1134 (2012).

48. Miller, R., Plessow, F., Kirschbaum, C. & Stalder, T. Classification criteria for distinguishing cortisol responders from nonresponders to psychosocial stress: evaluation of salivary cortisol pulse detection in panel designs. Psychosom. Med. 75, 832–840 (2013).

49. Hahn, A. et al. Reconfiguration of functional brain networks and metabolic cost converge during task performance. eLife 9, e52443 (2020).

50. Alves, P. N. et al. An improved neuroanatomical model of the default-mode network reconciles previous neuroimaging and neuropathological findings. *Commun*. Biol. 2, 370 (2019).

51. Klug, S. et al. Learning induces coordinated neuronal plasticity of metabolic demands and functional brain networks. *Commun*. Biol. 5, 428 (2022).

52. Godbersen, G. M. et al. Non-invasive assessment of stimulation-specific changes in cerebral glucose metabolism with functional PET. Eur. J. Nucl. Med. Mol. Imaging 51, 2283–2292 (2024).

53. Riedl, V. et al. Metabolic connectivity mapping reveals effective connectivity in the resting human brain. Proc. Natl. Acad. Sci. U. S. A. 113, 428–433 (2016).

54. Sicorello, M. et al. The functional neurobiology of dispositions towards negative emotions. Nat. Commun. 17, 5622 (2026).

55. Berretz, G., Packheiser, J., Kumsta, R., Wolf, O. T. & Ocklenburg, S. The brain under stress-A systematic review and activation likelihood estimation meta-analysis of changes in BOLD signal associated with acute stress exposure. Neurosci. Biobehav. Rev. 124, 89–99 (2021).

56. Raichle, M. E. et al. A default mode of brain function. Proc. Natl. Acad. Sci. 98, 676–682 (2001).

57. Zhang, W. et al. Acute stress alters the ‘default’ brain processing. NeuroImage 189, 870–877 (2019).

58. Buckner, R. L., Andrews-Hanna, J. R. & Schacter, D. L. The brain’s default network: anatomy, function, and relevance to disease. Ann. N. Y. Acad. Sci. 1124, 1–38 (2008).

59. Fox, M. D. et al. The human brain is intrinsically organized into dynamic, anticorrelated functional networks. Proc. Natl. Acad. Sci. U. S. A. 102, 9673–9678 (2005).

60. King, S. et al. Characterising a stress-sensitive default mode network (DMN) deficit in major psychiatric disorders. *Commun*. Biol. 9, 603 (2026).

61. Dixon, M. L. et al. Frontoparietal and Default Mode Network Contributions to Self-Referential Processing in Social Anxiety Disorder. Cogn. Affect. Behav. Neurosci. 22, 187–198 (2022).

62. Gu, H., Hu, Y., Chen, X., He, Y. & Yang, Y. Regional excitation-inhibition balance predicts default-mode network deactivation via functional connectivity. NeuroImage 185, 388–397 (2019).

63. Buzsáki, G., Kaila, K. & Raichle, M. Inhibition and brain work. Neuron 56, 771–783 (2007).

64. Hu, Y., Chen, X., Gu, H. & Yang, Y. Resting-state glutamate and GABA concentrations predict task-induced deactivation in the default mode network. J. Neurosci. Off. J. Soc. Neurosci. 33, 18566–18573 (2013).

65. Popoli, M., Yan, Z., McEwen, B. & Sanacora, G. The stressed synapse: the impact of stress and glucocorticoids on glutamate transmission. Nat. Rev. Neurosci. 13, 22–37 (2011).

66. Pellerin, L. & Magistretti, P. J. Glutamate uptake into astrocytes stimulates aerobic glycolysis: a mechanism coupling neuronal activity to glucose utilization. Proc. Natl. Acad. Sci. U. S. A. 91, 10625–10629 (1994).

67. Cole, M. W. et al. Multi-task connectivity reveals flexible hubs for adaptive task control. Nat. Neurosci. 16, 1348–1355 (2013).

68. Wang, R., Zhen, S., Zhou, C. & Yu, R. Acute stress promotes brain network integration and reduces state transition variability. Proc. Natl. Acad. Sci. U. S. A. 119, e2204144119 (2022).

69. Hermans, E. J., Henckens, M. J. A. G., Joëls, M. & Fernández, G. Dynamic adaptation of large-scale brain networks in response to acute stressors. Trends Neurosci. 37, 304–314 (2014).

70. Baker, J. T. et al. Functional connectomics of affective and psychotic pathology. Proc. Natl. Acad. Sci. U. S. A. 116, 9050–9059 (2019).

71. Yan, C.-G. et al. Reduced default mode network functional connectivity in patients with recurrent major depressive disorder. Proc. Natl. Acad. Sci. U. S. A. 116, 9078–9083 (2019).

72. Ironside, M. et al. Reductions in rostral anterior cingulate GABA are associated with stress circuitry in females with major depression: a multimodal imaging investigation. Neuropsychopharmacol. Off. Publ. Am. Coll. Neuropsychopharmacol. 46, 2188–2196 (2021).

73. Cunningham, S. et al. Cortisol reactivity to stress predicts behavioral responsivity to reward moderation by sex, depression, and anhedonia. J. Affect. Disord. 293, 1–8 (2021).

74. Menon, V. The Triple Network Model, Insight, and Large-Scale Brain Organization in Autism. Biol. Psychiatry 84, 236–238 (2018).

75. Menon, V. & Uddin, L. Q. Saliency, switching, attention and control: a network model of insula function. Brain Struct. Funct. 214, 655–667 (2010).

76. Menon, V. Large-scale brain networks and psychopathology: a unifying triple network model. Trends Cogn. Sci. 15, 483–506 (2011).

77. Craig, A. D. B. How do you feel--now? The anterior insula and human awareness. Nat. Rev. Neurosci. 10, 59–70 (2009).

78. Craig, A. D. Interoception: the sense of the physiological condition of the body. Curr. Opin. Neurobiol. 13, 500–505 (2003).

79. McEwen, B. S. et al. Mechanisms of stress in the brain. Nat. Neurosci. 18, 1353–1363 (2015).

80. Southwick, S. M. & Charney, D. S. The science of resilience: implications for the prevention and treatment of depression. Science 338, 79–82 (2012).

81. Charney, D. S. Psychobiological mechanisms of resilience and vulnerability: implications for successful adaptation to extreme stress. Am. J. Psychiatry 161, 195–216 (2004).

82. Vanderwal, T., Kelly, C., Eilbott, J., Mayes, L. C. & Castellanos, F. X. Inscapes: A movie paradigm to improve compliance in functional magnetic resonance imaging. NeuroImage 122, 222–232 (2015).

83. Soliman, A. et al. Limbic response to psychosocial stress in schizotypy: a functional magnetic resonance imaging study. Schizophr. Res. 131, 184–191 (2011).

84. Connor, K. M. & Davidson, J. R. T. Development of a new resilience scale: the Connor-Davidson Resilience Scale (CD-RISC). Depress. Anxiety 18, 76–82 (2003).

85. Wright, K. L. et al. Clinical evaluation of CAIPIRINHA: comparison against a GRAPPA standard. J. Magn. Reson. Imaging JMRI 39, 189–194 (2014).

86. Burgos, N. et al. Attenuation correction synthesis for hybrid PET-MR scanners: application to brain studies. IEEE Trans. Med. Imaging 33, 2332–2341 (2014).

87. Milz, C. et al. Open-access template and database approaches for pseudo-CT generation in brain PET/MRI attenuation correction. 2025.12.22.695942 Preprint at 10.64898/2025.12.22.695942 (2025).

88. Hahn, A. et al. A Unified Approach for Identifying PET-based Neuronal Activation and Molecular Connectivity with the functional PET toolbox. 2024.11.13.623377 Preprint at 10.1101/2024.11.13.623377 (2024).

89. Afyouni, S. & Nichols, T. E. Insight and inference for DVARS. NeuroImage 172, 291–312 (2018).

90. Phạm, D. Đ., McDonald, D. J., Ding, L., Nebel, M. B. & Mejia, A. F. Less is more: balancing noise reduction and data retention in fMRI with data-driven scrubbing. NeuroImage 270, 119972 (2023).

91. Sun, F. T., Miller, L. M. & D’Esposito, M. Measuring interregional functional connectivity using coherence and partial coherence analyses of fMRI data. NeuroImage 21, 647–658 (2004).

92. Attwell, D. & Laughlin, S. B. An energy budget for signaling in the grey matter of the brain. J. Cereb. Blood Flow Metab. Off. J. Int. Soc. Cereb. Blood Flow Metab. 21, 1133–1145 (2001).

93. Klein, A. & Tourville, J. 101 labeled brain images and a consistent human cortical labeling protocol. Front. Neurosci. 6, 171 (2012).

94. Hahn, A. et al. Escitalopram enhances the association of serotonin-1A autoreceptors to heteroreceptors in anxiety disorders. J. Neurosci. Off. J. Soc. Neurosci. 30, 14482–14489 (2010).

