## supplementary figure 1a for "Reconfiguration of default mode metabolic demands and network organization during acute stress"

### Supplementary Material

#### Supplementary results

As sex differences in stress processing have been reported in previous work,[1,2] we specifically tested for differences between women and men. However, no significant differences (all p-values uncorrected) were observed with respect to the number of stress responders based on ACTH levels ($\chi^2$=1.5, p=0.23), stress-specific glucose metabolism in the PCC (t=1.8, p=0.08) and ACC (t=0.9, p=0.9) nor for BOLD signal changes in the PCC (t=1.4, p=0.15).

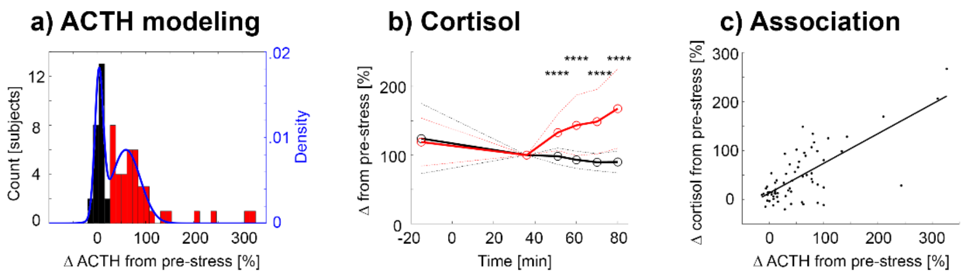


**Supplementary Figure S1: Additional results on the endocrine stress response.** a) Identification of stress responders and non-responders by modelling change in ACTH levels with a Gaussian mixture model (two Gaussians, values above 142% change (i.e., outliers as defined by boxplot) were not used for the model). The cut-off was determined as 25.8%, yielding 62.1% of participants as stress responders. b) Cortisol profiles using a conventional threshold of 15% increase in cortisol levels from baseline[3] to identify stress responders and non-responders, yielding 63.6% responders. Comparing the stress response rate with cortisol levels here to that obtained when using ACTH levels (Figure 2) indicated high concordance (chi-square test p < 0.0001). Separation between responders and non-responders was also similar when using changes in ACTH or cortisol levels (two sample t-test, both p < 0.0001). c) A moderate correlation of percent change between the two different hormone parameters was observed (Spearman's rho = 0.58, p < 0.0001). red: stress responders, black: stress non-responders.

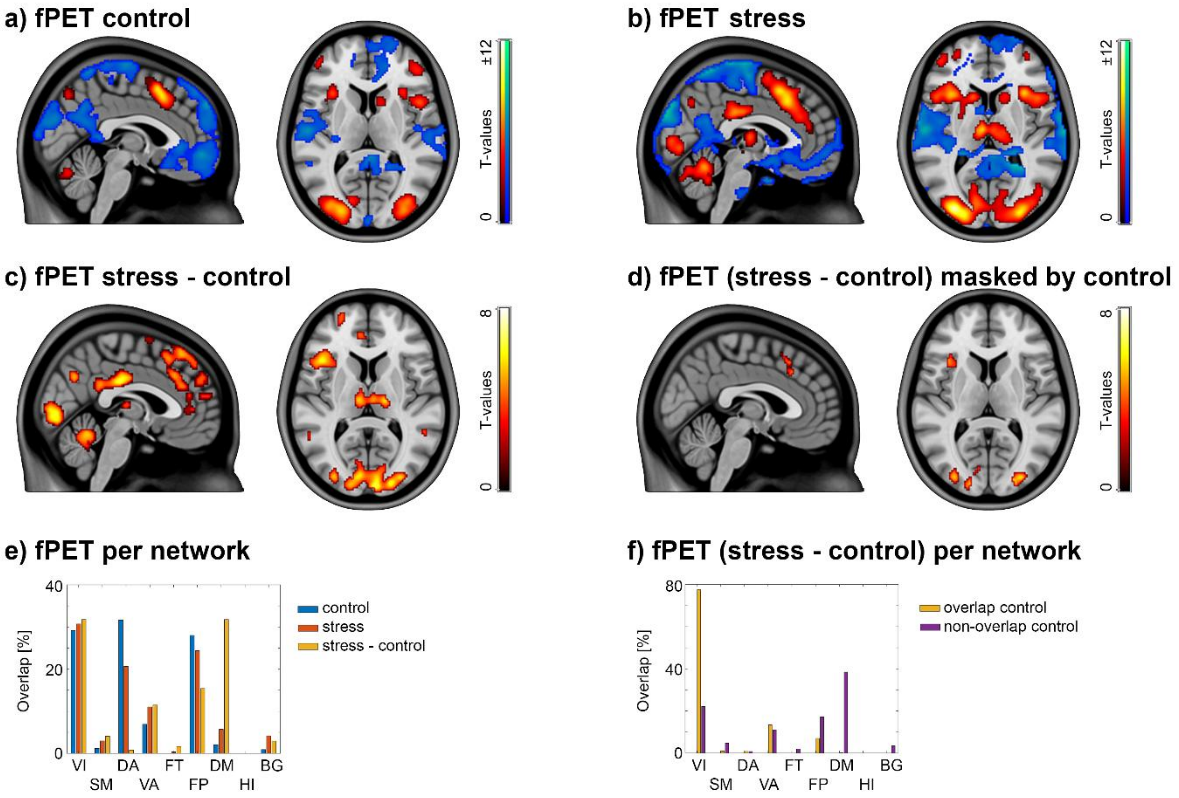


**Supplementary Figure S2: Different [18F]FDG fPET statistical contrasts.** a) Changes in CMRGlu for the control condition as compared to baseline radiotracer uptake. b) Changes in CMRGlu for the stress condition as compared to baseline radiotracer uptake. c) Significantly higher CMRGlu during stress vs. control. d) Same as c, but differences are masked by effects in a. e) Percentage overlap of significant effects in the respective brain networks for the different contrasts. f) The overlap of the effect stress – control per network is further separated for those voxels that overlap with the control condition (yellow) vs. those that do not overlap (purple). Together, these effects demonstrate the specific increase of CMRGlu in the regions of the DMN, which are recruited de novo during stress. All images (a - d) are shown at $p<0.05$ FWE corrected cluster level following $p<0.001$ uncorrected voxel level. Panel c and yellow bars in panel e are replicated from figure 3.

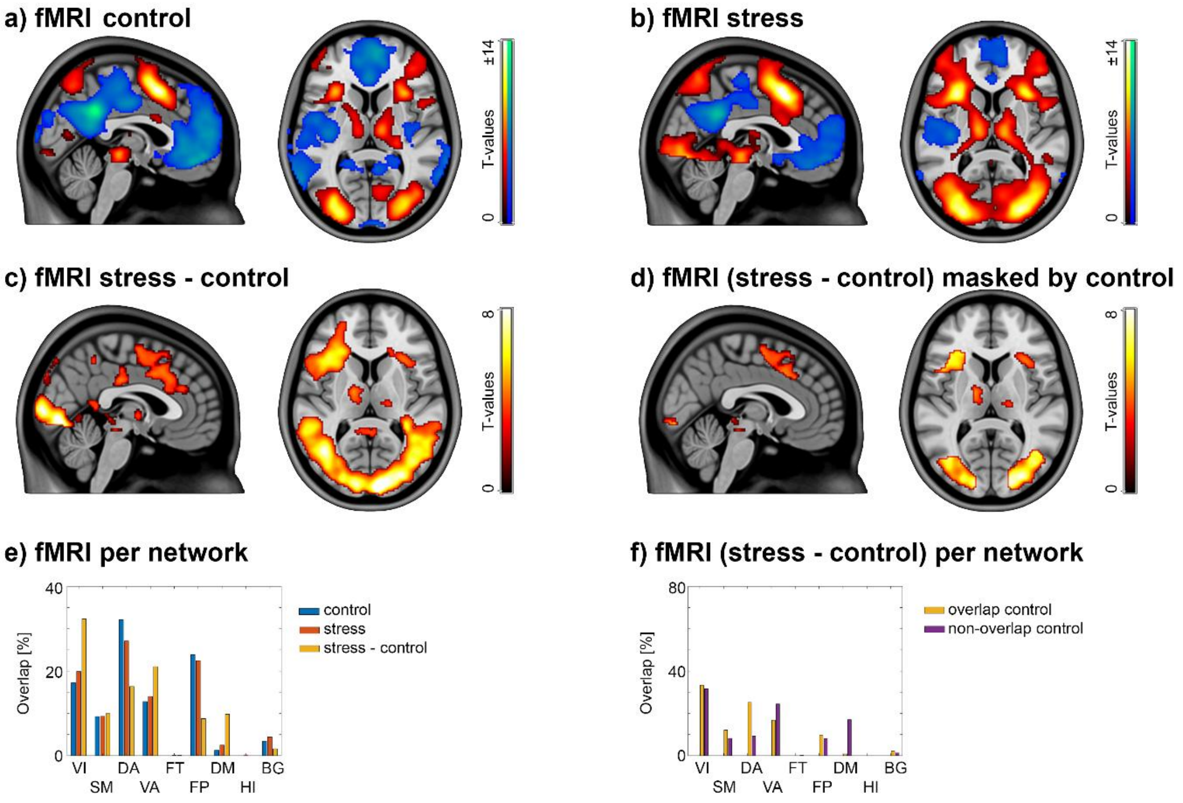


**Supplementary figure S3: Different BOLD fMRI statistical contrasts.** a) Changes in the BOLD signal for the control condition as compared to baseline. b) Changes in the BOLD signal for the stress condition as compared to baseline. c) Significantly higher changes in the BOLD signal during stress vs. control. d) Same as c, but differences are masked by effects in a. e) Percentage overlap of significant effects in the respective brain networks for the different contrasts. f) The overlap of the effect stress – control per network is further separated for those voxels that overlap with the control condition (yellow) vs. those that do not overlap (purple). Together, these effects demonstrate that BOLD signal decreases of the control condition in the DMN are alleviated during stress. All images (a - d) are shown at $p<0.05$ FWE corrected cluster level following $p<0.001$ uncorrected voxel level. Panel c and yellow bars in panel e are replicated from figure 3.

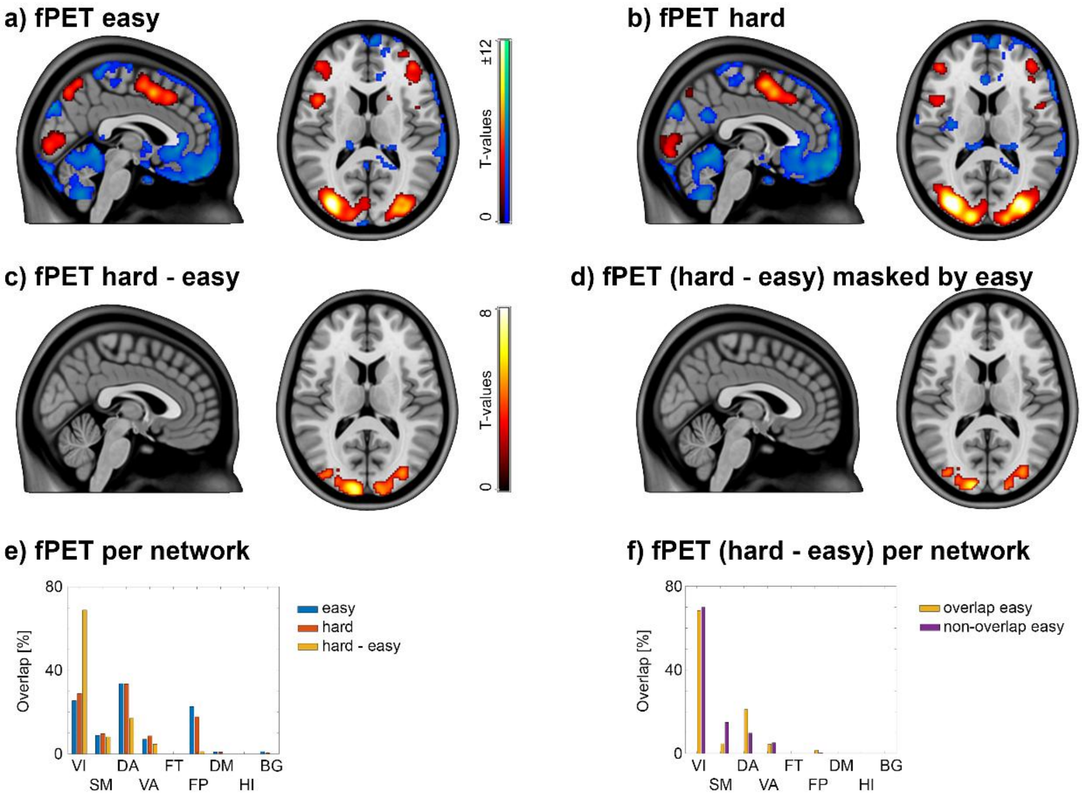


**Supplementary Figure S4: Different [18F]FDG fPET statistical contrasts of a cognitive challenge task that does not involve psychosocial stress (Tetris).** a) Changes in CMRGlu for the easy task condition as compared to baseline radiotracer uptake. b) Changes in CMRGlu for the hard task condition as compared to baseline radiotracer uptake. c) Significantly higher CMRGlu during hard vs. easy. d) Same as c, but differences are masked by effects in a. e) Percentage overlap of significant effects in the respective brain networks for the different contrasts. f) The overlap of the effect hard – easy per network is further separated for those voxels that overlap with the easy task condition (yellow) vs. those that do not overlap (purple). Together, these effects demonstrate the majority of increases in CMRGlu emerge in the visual cortex and most of them overlap with the control condition. All images (a - d) are shown at $p<0.05$ FWE corrected cluster level following $p<0.001$ uncorrected voxel level. All data were replicated from previous work.[4]

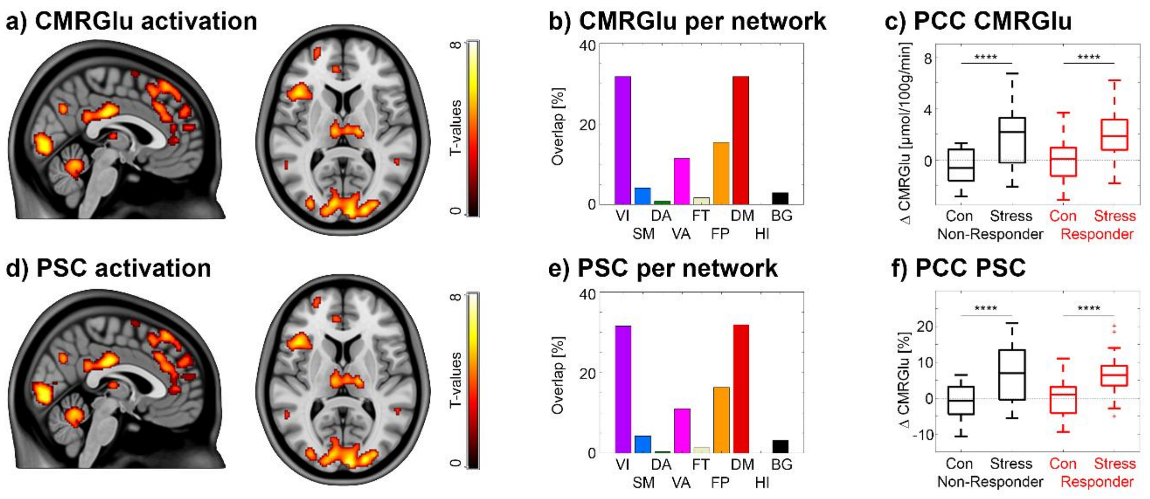


**Supplementary figure S5: Comparison of [$^{18}$F]FDG fPET results** when using CMRGlu and PSC as outcome parameters. a-c) Main effect of condition (stress vs. control) with CMRGlu, replicated from figure 3. d-f) The same statistics used in a-c, but with PSC as outcome parameter. All images (a, d) are shown at p<0.05 FWE corrected cluster level following p<0.001 uncorrected voxel level. ****$p_{Bonf} < 0.0001$.

a) ACC CMRGlu

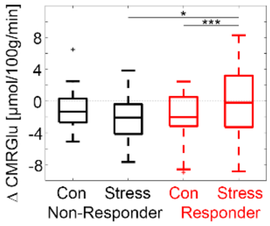


**Supplementary figure S6: Glucose metabolism of the ACC** as a function of stress responsiveness when defined with the established threshold of 15% change in cortisol levels[3] (interaction effect as in figure 4). Stress responder showed higher CMRGlu when compared to the control condition ($p<0.001$) and to non-responder during stress ($p<0.05$).

#### References:


1. Dong, D. *et al.* Sex-specific neural responses to acute psychosocial stress in depression. *Transl. Psychiatry* **12**, 2 (2022).
2. Kudielka, B. M. & Kirschbaum, C. Sex differences in HPA axis responses to stress: a review. *Biol. Psychol.* **69**, 113–132 (2005).
3. Miller, R., Plessow, F., Kirschbaum, C. & Stalder, T. Classification criteria for distinguishing cortisol responders from nonresponders to psychosocial stress: evaluation of salivary cortisol pulse detection in panel designs. *Psychosom. Med.* **75**, 832–840 (2013).
4. Hahn, A. *et al.* Reconfiguration of functional brain networks and metabolic cost converge during task performance. *eLife* **9**, e52443 (2020).
